# Does pre-speech auditory modulation reflect processes related to feedback monitoring or speech movement planning?

**DOI:** 10.1101/2024.07.13.603344

**Authors:** Joanne Jingwen Li, Ayoub Daliri, Kwang S. Kim, Ludo Max

## Abstract

Previous studies have revealed that auditory processing is modulated during the planning phase immediately prior to speech onset. To date, the functional relevance of this pre-speech auditory modulation (PSAM) remains unknown. Here, we investigated whether PSAM reflects neuronal processes that are associated with preparing auditory cortex for optimized feedback monitoring as reflected in online speech corrections. Combining electroencephalographic PSAM data from a previous data set with new acoustic measures of the same participants’ speech, we asked whether individual speakers’ extent of PSAM is correlated with the implementation of within-vowel articulatory adjustments during /b/-vowel-/d/ word productions. Online articulatory adjustments were quantified as the extent of change in inter-trial formant variability from vowel onset to vowel midpoint (a phenomenon known as *centering*). This approach allowed us to also consider inter-trial variability in formant production and its possible relation to PSAM at vowel onset and midpoint separately. Results showed that inter-trial formant variability was significantly smaller at vowel midpoint than at vowel onset. PSAM was not significantly correlated with this amount of change in variability as an index of within-vowel adjustments. Surprisingly, PSAM was negatively correlated with inter-trial formant variability not only in the middle but also at the very onset of the vowels. Thus, speakers with more PSAM produced formants that were already less variable at vowel onset. Findings suggest that PSAM may reflect processes that influence speech acoustics as early as vowel onset and, thus, that are directly involved in motor command preparation (feedforward control) rather than output monitoring (feedback control).

## 1. Introduction

Previous studies suggest that auditory processing is modulated during speaking. For example, auditory cortex has been found to show an attenuated response to self-generated speech compared with externally generated speech, a phenomenon called speaking-induced suppression (SIS) [1–6]. SIS has been interpreted as a partial neural cancellation in response to sensory feedback that matches the predicted sensory consequences of motor commands [7–10].

A series of electroencephalography (EEG) studies from our own laboratory have revealed that auditory processing in typical speakers is already modulated during speech movement planning prior to speech onset. Such pre-speech auditory modulation (PSAM) is observed as a reduction in amplitude of the N1 component in Long Latency Auditory Evoked Potentials (LLAEP) in response to auditory stimuli presented immediately prior to speaking as compared with a control condition without speaking [11–16]. We previously hypothesized that PSAM may reflect a preparation of the auditory cortex associated with adjusting the facilitation-suppression balance of auditory neuronal populations [17] to optimize subsequent feedback monitoring while speaking [12–16]. However, we recently found evidence suggesting that this may not be the case. Specifically, using psychophysics methods, Wang, Ali, and Max [18] found that, in the time window where PSAM is observed, speakers’ ability to detect a small deviation in vowel formant frequencies is diminished rather than enhanced — suggesting a general attenuation of auditory processing during speech planning and not a change to a state that is more optimized for acoustic error detection.

We therefore addressed in the current study more directly whether PSAM reflects neuronal processes associated with preparing auditory cortex for optimized feedback monitoring by assessing its relation to online speech corrections. We combined EEG PSAM data and acoustic speech data from two separate tasks completed by the same participants in a previous study [15]. First, we assessed whether individual speakers’ extent of PSAM is correlated with the extent of *change* in inter-trial formant variability from vowel onset to vowel midpoint. This acoustics-based measure is known as *centering* and quantifies how much change occurred in vowel dispersion from onset to midpoint relative to the speaker’s own median onset and midpoint for that vowel [6, 19–21]. Thus, the measure quantifies the extent to which produced vowels moved closer to the median production from onset to midpoint, and it can be used as an index of within-vowel articulatory adjustments. Second, extracting formant frequency measures at both vowel onset and midpoint also allowed us to assess the relationship between the extent of PSAM and inter-trial variability in formant production *separately* at vowel onset and vowel midpoint.

Overall, the underlying premise was that variability at vowel onset indicates the robustness of feedforward planning whereas variability at vowel midpoint reflects both the planned output and the implementation of any online feedback-based corrections.

## 2. Methods

### 2.1. Participants

The included data were obtained from twenty-four adults between 19 and 45 years of age. This overall group included 12 typical individuals with no prior or current speech difficulties and 12 speakers who stutter (a population known to have reduced PSAM, thus providing the opportunity to study the experimental question of interest over a wider range of PSAM values). All participants were native speakers of American English, right-handed, and unaware of the study’s purpose. They had passed a pure-tone hearing screening (20 dB HL for octave frequencies of 250-4000 Hz) for both ears separately, reported to have no psychological, neurological, or communication disorders (except for stuttering among the participants who stutter), and no medication use that could affect sensorimotor functioning. Details about the individual participants are available in [15].

### 2.2. Data

The analyzed data include acoustic measures of the first and second formant frequency (F1, F2) in monosyllabic words produced during the baseline phase of the speech adaptation task in [15] and EEG-based measures of PSAM recorded during the separate PSAM task in the same study. The two tasks were conducted either on the same day or 3–15 days apart.

#### 2.2.1. Acoustic speech data

For the present study, we included acoustic measures of F1 and F2 that had been extracted from the participants’ repeated productions of three consonant-vowel-consonant (CVC) words *during the unperturbed baseline phase* of *the first iteration* of a speech adaptation task. Thus, although participants were wearing insert earphones (see below), all productions included here were recorded prior to the introduction of any auditory feedback perturbation and without any participant awareness that a perturbation would occur later.

The three words produced during the task were “bed” (/bεd/), “bud” (/bʌd/), and “baud” (/bɔd/). These words occurred in randomized order in each block of 3 trials. The data available for the present study (i.e., only the baseline phase of the task) included 10 blocks, yielding 30 trials for each of the 24 participants.

Participants produced the words while seated in a sound-attenuated booth, wearing insert earphones (ER-3A, Etymotic Research Inc., Grove Village, IL), and facing a computer monitor approximately 1.5 m away. Participants were instructed to produce the target words, presented one at a time on the monitor, while maintaining their volume at a consistent level (65-71 dB SPL) with the assistance of color-coded feedback. A microphone (WL185, Shure Incorporated, Niles, IL) was placed 15 cm in front of the participant’s mouth to record the acoustic speech output. This speech signal was amplified (DPS II, ART ProAudio, Niagara Falls, NY) and sent through a speech processor (VoiceOne, TC-Helicon, Victoria, BC, Canada) but, as mentioned, only utterances produced prior to the introduction of altered feedback were included for the present study. The speech signal was routed to a headphone amplifier (S-phone, Samson Technologies Corp., Syosset, NY) and played back to the participants through the insert earphones with a latency of only 11 ms [22]. Before each session started, the volume of the speech signal was calibrated such that a speech sound of 68 dB SPL recorded at the microphone corresponded to an output of 73 dB SPL in the insert earphones.

#### 2.2.2. EEG PSAM data

Participants’ PSAM data are those reported in [15]. The two conditions used to derive PSAM measures were a speaking condition and a silent reading condition, each consisting of three blocks of 90 trials. In the speaking condition, each trial started with the presentation of a target word in white characters on a computer monitor with black background. After 600 ms, the word color turned to green, and this served as a *go*-signal for the participants to read the word out loud. In 40% of the trials (tone trials) within each block, a 1 kHz, 40ms duration pure tone was presented at 75 dB SPL in the insert earphones 400 ms after the onset of presentation of the target word (Figure 1.A). The remaining trials were no-tone trials. The interval between successive trials was randomly selected from five possible values (1500, 2000, 2500, 3000, 3500 ms). The silent reading condition was identical to the speaking condition except that participants were instructed to read the words silently without any orofacial movements.

**Figure 1.**
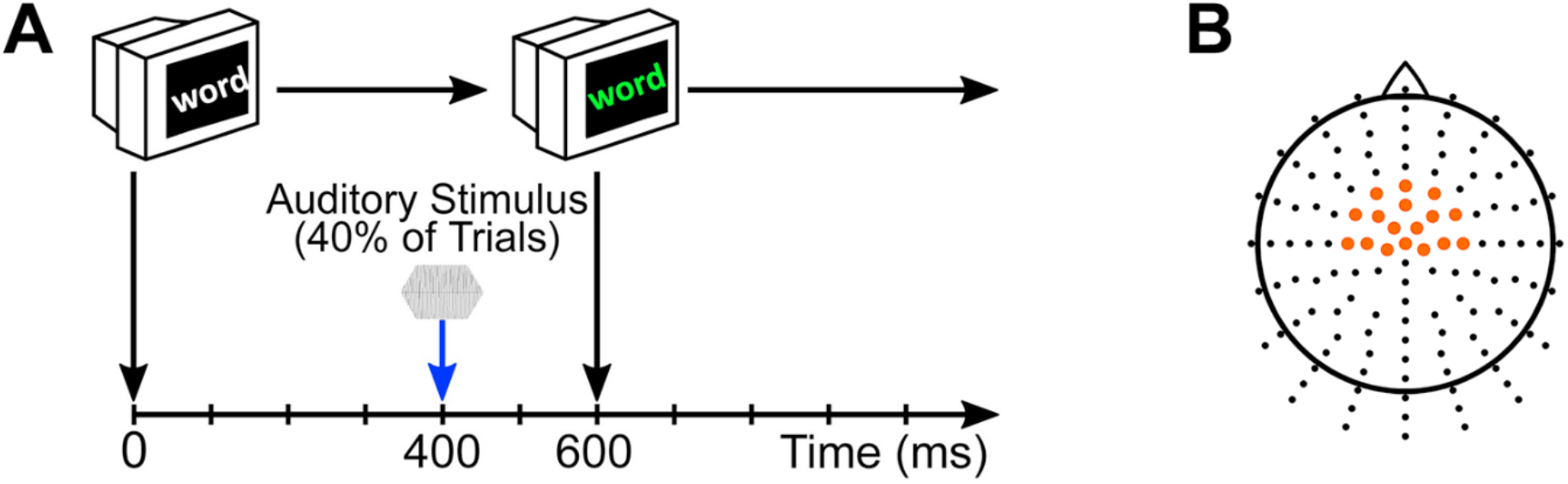
(A) Timeline of visual and auditory stimulus presentation for electroencephalographic (EEG) recordings of pre-speech auditory modulation (PSAM). (B) Electrode channels averaged for PSAM analysis.

EEG recordings (Active Two, BioSemi, Amsterdam, The Netherlands) were made from 128 electrodes in an elastic cap with a layout extending the international 10-10 system [23, 24]. The data used here were based on a subset of 17 data channels in the frontocentral region (Figure 1.B) where PSAM had been found to be prominent [13]. The signals from two electrodes positioned over the left and right mastoids were used to calculate an averaged mastoid reference during offline analysis. A bipolar EOG signal, recorded using electrodes placed below and next to the left eye, allowed for the rejection of blinks and eye movements. Upper and lower lip movements were detected by recording monopolar surface EMG signals using four electrodes positioned over the right and left parts of orbicularis oris superior and inferior.

### 2.3. Data extraction and analysis^1^

#### 2.3.1. Acoustic variability

Measurements of the acoustic data were completed in MATLAB (The Mathworks, Natick, MA) using custom written scripts. The first two formant frequencies (F1, F2) of vowels in the participants’ productions were automatically extracted using formant tracking algorithms available in Praat [25]. We visually inspected spectrograms with overlaid formant tracks to ensure accuracy. When needed, Praat’s formant tracking parameters were adjusted. We used the spectrograms also to manually label onset and offset of the vowel for each production.

The formants at vowel onset and midpoint were calculated as the average F1 and F2 across time windows corresponding to the first 20% and middle 20% of the vowels, respectively. All formant values were converted from the original hertz units to the mel scale. For vowel onset and midpoint separately, we calculated inter-trial formant variability by measuring the Euclidean distance (in F1-F2 coordinates) from each token to the median of the data on a per-word basis (i.e., separately for “bed”, “bud”, “baud”) and then averaging across the three target vowels. We calculated the change in inter-trial formant variability by subtracting the variability at vowel midpoint from that at vowel onset, which is consistent with *centering* analyses in the literature [6, 20].

#### 2.3.2. PSAM

EEG signals were re-referenced to the average mastoid signals and filtered using a 50 Hz low-pass filter (Kaiser windowed sinc FIR filter; deviation .005; transition bandwidth 1 Hz). The filter was applied both forward and backward to maintain zero-phase shift [26]. The remaining signals were segmented into epochs from –100 ms to 400 ms relative to the auditory stimulus onset in the tone trials and the same timepoint in the no-tone trials. The epochs were then baseline-corrected by subtracting the average amplitude across the –100 ms time window from the entire epoch. Next, we visually inspected the epochs and excluded those with amplitudes exceeding ±100 μV and those showing artifacts related to excessive muscle activity, muscle activity prior to the go-signal, blinking, and eye movements.

To isolate neural activity specifically evoked by the auditory stimuli, we calculated separate averages for the epochs for all tone trials and the epochs for all no-tone trials. The average signal of the tone trials reflects combined brain activity in response to the auditory stimuli and that associated with non-auditory processes (e.g., motor, cognitive, etc.) whereas the average signal of the no-tone trials only reflects non-auditory processes. Accordingly, we subtracted for each subject the average signal for no-tone trials from that for tone trials to obtain the best estimate of the auditory activity. A low-pass filter at 15 Hz (Kaiser windowed sinc FIR filter; deviation: .005; transition bandwidth: 1 Hz) was applied to the signals to obtain each channel’s final LLAEP [13, 27–30]. Lastly, for each subject, we averaged the LLAEPs from 17 channels located in the frontocentral region (illustrated in Figure 1.B). The amplitude of the N1 component (defined as the largest negative peak between 70 and 130 ms after onset of the auditory stimulus) in this evoked potential was calculated, and PSAM was measured as the difference in N1 amplitude between the speaking and silent reading conditions. Positive PSAM values indicate that the negative N1 amplitude was reduced in the speaking condition relative to the silent reading condition.

#### 2.3.3. Statistical analyses

All statistical analyses were completed in R version 4.3.2 base packages [31] unless otherwise noted. First, to determine whether there was a significant reduction in inter-trial formant variability from vowel onset to vowel midpoint, we conducted a paired-samples *t*-test with all participants’ formant variability data from those two time points. In addition, Pearson correlation analyses were used to examine whether a speaker’s change in formant variability from vowel onset to midpoint is related to their amount of PSAM. We also examined the association between PSAM and variability at vowel onset and midpoint separately. Lastly, we compared the latter two correlation coefficients with the *r.test* function in the “psych” package, implementing a Williams’ *t* test for dependent correlations with a shared variable [32].

## 3. Results

Inter-trial formant variability showed a statistically significant decrease from vowel onset to vowel midpoint (*t(23) = -14.354, p < .001, d = -2.930*) (Figure. 2.A). Across speakers, this *change* in formant variability from vowel onset to midpoint was not statistically significantly correlated with the amount of PSAM (*r = 0.128, p = .552*) (Figure. 2.B). However, PSAM was statistically significantly correlated with inter-trial formant variability at *both* vowel onset (r *= - 0.453, p = .026*) (Figure. 2.C) and vowel midpoint (*r = -0.669, p < .001*) (Figure. 2.D). At both vowel onset and midpoint, individuals who showed a greater amount of PSAM produced the vowels with less inter-trial variability. The two separate correlation coefficients obtained for vowel onset and vowel midpoint were not statistically significantly different (*t = 1.538, p = .139*).

**Figure 2.**
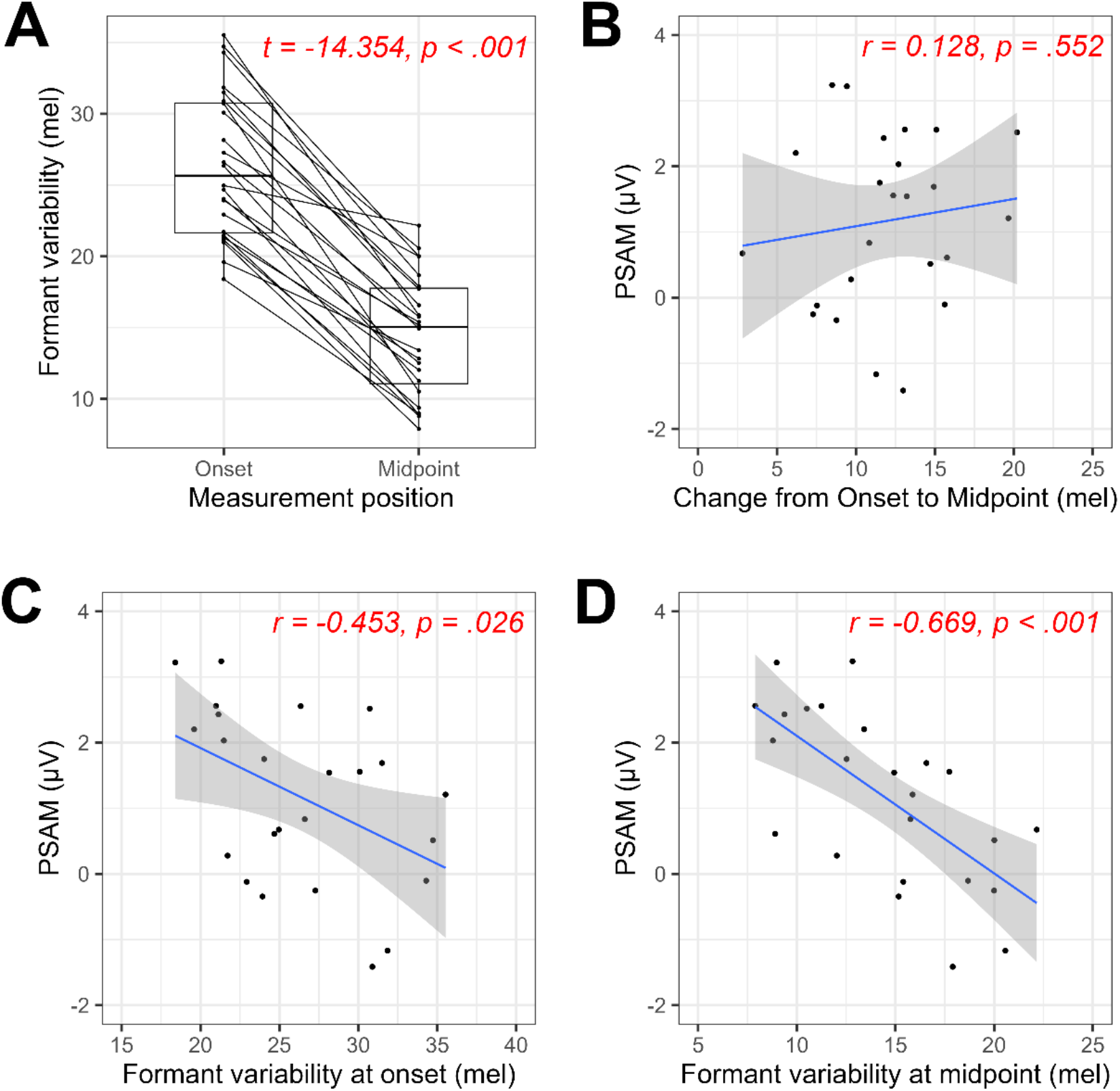
(A) Change in inter-trial formant variability from vowel onset to midpoint. (B) Correlation between PSAM and change in inter-trial formant variability from vowel onset to midpoint. (C) Correlation between PSAM and inter-trial formant variability at vowel onset. (D) Correlation between PSAM and inter-trial formant variability at vowel midpoint.

## 4. Discussion

Our previous work showed that speech planning prior to movement onset is associated with a modulation of auditory processing [12–16]. With regard to the functional relevance of this PSAM phenomenon, we previously hypothesized that it may reflect neural processes related to optimizing auditory cortex’s sensitivity and frequency specificity for subsequent auditory feedback monitoring [16]. However, our more recent results from a psychophysics study did not corroborate this hypothesis [18]. In the present study, we therefore aimed to investigate more directly whether a speaker’s extent of PSAM during the planning phase immediately prior to speech onset (a measure derived from EEG data) correlates with the extent of online corrections in the formant frequencies of their produced utterances (an acoustic measure reflecting articulatory behavior). Given that this approach required formant frequency measures to be made at both vowel onset and vowel midpoint, we were also able to assess the relationship between extent of PSAM and behavioral inter-trial variability at vowel onset and vowel midpoint separately. The latter analysis provided an opportunity to test the alternative hypothesis that PSAM relates not to fine-tuning auditory cortex for feedback-based corrections during the production but to the involvement of auditory cortex in feedforward movement planning itself, a process already completed by the time of speech onset.

First, we found that inter-trial formant variability decreased significantly from vowel onset to midpoint, consistent with previous reports of within-vowel *centering* [6, 20, 21]. Although Niziolek and colleagues [6, 19] have interpreted their data as indicating that the involved articulatory adjustments are at least partially driven by auditory feedback, it should be noted that one study found that neither typical speakers nor cerebellar ataxia patients showed a difference in the extent of centering when auditory feedback was or was not available [21]. Thus, the involvement of corrections specifically for *auditory* error remains ambiguous. Most importantly, as our second finding, the present study revealed no significant correlation between speakers’ amount of PSAM and their extent of within-vowel centering. In other words, regardless of whether or not the within-vowel articulatory adjustments are partially attributable to auditory error correction mechanisms, the specific adjustments investigated here do not relate to the neural processes reflected in PSAM.

As a third finding, our results did reveal a significant negative correlation between PSAM and inter-trial formant variability. Speakers who exhibited more PSAM during speech movement planning produced vowels with less inter-trial formant variability. Surprisingly, this correlation was statistically significant not only at vowel midpoint but even as early as vowel onset. In fact, the strength of the correlation was not statistically significantly different for PSAM and formant variability at vowel onset as compared with PSAM and formant variability at vowel midpoint.

Given that this finding is of a correlational nature, one possibility is that both PSAM and intertrial formant variability are driven by the common influence of a third variable. As an example, it is theoretically possible that a variable such as the size of speakers’ auditory goals in formant space would influence both the preparation for precise feedback monitoring and the amount of inter-trial variability. If so, it would be reasonable to expect that larger auditory goals would lead to both more inter-trial variability and less preparation for auditory monitoring, consistent with the direction of the variability-PSAM correlation observed in our data. Indeed, it has previously been suggested that greater formant-space dispersion of multiple productions of the same vowel indicates that the speaker has larger auditory goals in this space, and that auditory feedback monitoring rejects/accepts individual productions based on the speaker’s goal size [33]. However, according to this same perspective, speakers with larger auditory goals should also show less auditory-motor adaptation in response to formant-shift perturbations in the real-time auditory feedback [33], and our own prior results do not provide support for a correlation between PSAM and the extent of auditory-motor adaptation [15]. Thus, at the present time, it is doubtful that our new finding of a correlation between PSAM and inter-trial formant variability as early as vowel onset is best explained by a mediating role for the size of speakers’ auditory goals. Of course, it cannot be ruled out that other variables may play such a role and this possibility will need to be investigated in future studies.

A second possible interpretation of our results is that the neural processes reflected in PSAM are not, as previously hypothesized, involved in preparing auditory cortex for optimized feedback monitoring but, instead, are directly involved in planning the *feedforward* motor commands for the subsequent speech movements. In combination, our findings of (a) a lack of correlation between PSAM and within-vowel changes, (b) the observation of a correlation between PSAM and formant variability at both vowel onset and midpoint, and (c) the absence of a statistical difference between the correlations at those two timepoints do indeed suggest that the effects of this auditory neural activity may already be fully established near the time of speech onset (recall that vowel onset measures were made for /b/-vowel-/d/ words). Thus, the reduced *sensory* response to probe tones during speech movement planning (i.e., PSAM; [11–13, 15] and the associate reduced *perceptual* discrimination of formant frequencies at the same timepoint [18] may be due to the simultaneous involvement of at least some auditory neuronal populations in bidirectional communication between pre-motor and sensory brain regions as part of *predictive* processes (i.e., without sensory input) that underlie the actual selection of feedforward motor commands.

The notion that the central nervous system generates sensory predictions by means of an internal forward model is a key concept in contemporary models of human sensorimotor control across effector systems (e.g., [34–40]). Typically, these perspectives postulate sensory predictions in the context of state estimation during ongoing movements, in particular as one component of an observer system integrating the predicted sensory state (based on efference copy and the forward model) and lagging sensory information about the actual state. However, besides this role in making faster online corrections and avoiding instabilities associated with feedback delays, sensory prediction may also play an important role in selecting the most optimal motor commands during movement planning.

One line of evidence strongly supporting this idea is found in a long history of studies documenting activation of relevant sensory cortical regions prior to movement onset in both humans and non-human primates [41–47]. For example, electrocorticography recordings from human participants during the preparation for finger movements showed that activation of primary somatosensory cortex (S1) immediately followed activation of premotor cortex (PM) and occurred before activation of primary motor cortex (M1); thus, sequential activation followed the order PM-S1-M1 [46]. Similarly, prior to speech onset, coherent neural activity has been observed between inferior frontal gyrus (which includes Broca’s area) and auditory cortex [47]. A second line of evidence supporting the involvement of sensory regions in movement planning comes from a number of recent studies on sensorimotor learning. First, in a finger force production task with varying force profile targets, changes in cortical excitability throughout learning occurred earlier in S1 than in M1 [48], suggesting that the earliest changes in movement planning may have depended primarily on changes in somatosensory cortex. Second, disrupting S1 with theta-burst transcranial magnetic stimulation (TMS) immediately after subjects adapted their reach movements to force field perturbations almost completely eliminated retention of learning [49], suggesting that S1 involvement is required for consolidation of newly learned sensorimotor transformations.

Given this extensive body of work confirming a critical role for (pre)motor-sensory interactions in movement planning, we propose that our present finding of an association between PSAM and inter-trial formant variability at vowel onset – in the absence of an association between PSAM and within-vowel changes – may reflect the direct involvement of auditory cortex in determining feedforward motor commands. Specifically, auditory cortex’s sensory response to the probe tones may be diminished during movement planning (i.e., PSAM occurs) due to recruitment of some of these neuronal populations for evaluating auditory predictions before movement initiation as part of the process of selecting the most optimal motor commands. This proposed interpretation of our results leads to the testable hypothesis that the articulatory movements of speakers with a greater amount of PSAM will also show a relatively greater contribution of feedforward control mechanisms. One logical next step, therefore, is to directly investigate the estimated weighting of feedforward versus feedback control mechanisms in the kinematics of unperturbed articulatory movements, for example, by quantifying to what degree movement endpoints are predictable from the initial movement kinematics [50, 51]. In the present study, we combined typical and stuttering speakers for the correlation analyses to be based on a wider spread of PSAM values (higher in typical speakers, lower in stuttering speakers), but such a future study with sufficiently large sample sizes would also allow for the potential association between PSAM and feedforward movement control to be examined within each group separately.

## Author contributions

**Joanne Jingwen Li**: Formal analysis; Visualization; Writing – original draft; Writing – review and editing. **Ayoub Daliri**: Methodology; Funding acquisition; Software; Investigation; Data curation; Formal analysis; Visualization; Writing – review and editing. **Kwang S. Kim**: Software; Formal analysis; Visualization; Writing – review and editing. **Ludo Max**: Conceptualization; Methodology; Funding acquisition; Project administration; Writing – original draft; Writing – review and editing.

## Footnotes

1 Data files are available at https://osf.io/2me8y/“

